# Generation of nanobodies targeting the human, transcobalamin-mediated vitamin B_12_ uptake route

**DOI:** 10.1101/2021.08.16.456495

**Authors:** Joël S. Bloch, Jeffrey M. Sequeira, Ana S. Ramírez, Edward V. Quadros, Kaspar P. Locher

**Author notes:** To whom correspondence should be addressed: Kaspar P. Locher, Otto-Stern-Weg 5, 8093 Zürich, Switzerland.

## Abstract

Cellular uptake of vitamin B_12_ in humans is mediated by the endocytosis of the B_12_ carrier protein transcobalamin (TC) via its cognate cell surface receptor TCblR (or CD320), encoded by the *CD320* gene(1). Because *CD320* expression is associated with the cell cycle and upregulated in highly proliferating cells such as cancer cells(2–4), this uptake route is a potential target for cancer therapy(5). We developed and characterized four camelid nanobodies that bind TC or the interface of the TC:TCblR complex with nanomolar affinities. We determined X-ray crystal structures of all four nanobodies in complex with TC:TCblR, which enabled us to map their binding sites. When conjugated to a toxin, three of these nanobodies are capable of inhibiting the growth of HEK293T cells and therefore have the potential to inhibit the growth of human cancer cells. We visualized the cellular binding and endocytic uptake of the most potent nanobody (TCNB4) using fluorescent light microscopy. The co-crystal structures of TC:TCblR with another nanobody (TCNB34) revealed novel features of the interface of TC and the LDLR-A1 domain of TCblR. Our findings rationalize the structural basis for a decrease in affinity of TC-B_12_ binding caused by the TCblR-Glu88 deletion mutant.

## Introduction

Because vitamin B_12_ is a cofactor of enzymes involved in recycling of folates for DNA synthesis(5), the need for B_12_ is elevated in highly proliferating cells such as pluripotent cancer cells(4). This correlates with the overexpression of the CD320 protein, the cellular receptor (TCblR, sometimes called CD320) of transcobalamin-B_12_ complex(4, 6). Targeting the cellular uptake route for vitamin B_12_ for potential cancer diagnosis or therapy has been described in previous studies(5), where strategies included the derivatization of vitamin B_12_(7) or the coupling of a receptor binding antibody to saporin(8, 9). In assays with cultured tissue, a saporin-conjugated antibody was able to inhibit a broad spectrum of cancer cell lines(9). Despite these efforts, no compound targeting the B_12_ uptake route is currently approved for clinical use(10). As an alternative way to exploit TCblR for potential cancer diagnosis or therapy, we have developed a set of nanobodies, which are heavy-chain-only antibody fragments that were originally derived from camelids and consist of a single IG domain. Nanobodies have found great popularity in basic science and biomedicine(11, 12) because they are highly stable, can be loaded with drugs, are less immunogenic than therapeutic antibodies, able to penetrate tissues, and can conveniently be produced at large scale in bacterial expression hosts including *E. coli*(13). Many nanobodies have been raised against cancer cell markers(12), and several companies have been funded to develop nanobody-based drugs. In 2019 Ablynx (Ghent, BEL) obtained the first FDA approval for a therapeutic nanobody while ten nanobodies from various companies were in clinical trials(14).

We hypothesized that a nanobody-drug conjugate targeting the cellular vitamin B_12_ uptake route might be advantageous over previous approaches and, therefore, could have potential use in cancer therapy and diagnosis. We generated a set of nanobodies targeting the TC-receptor complex, characterized them biophysically and using X-ray crystallography, and probed their uptake into human embryonic kidney (HEK293) cells.

## Materials and Methods

### Expression and purification of TC:TCblR

TC:TCblR complex was produced by co-transfection of the respective gene constructs into SF9 cells and purified as described previously(15).

### Nanobody selection, expression and purification

Nanobodies were generated, expressed and purified as described previously(16, 17). with the following modifications: For the panning TC:TCblR complex was immobilized on a streptavidin matrix via biotinylation of TCblR as described previously(15). Purified nanobodies were desalted into 20 mM Tris pH 7.5, 0.5 mM CaCl_2_ and 150 mM NaCl.

To generate nanobody-saporin conjugates, a C-terminal AviTag™ (GLNDIFEAQKIEWHE), preceding the His6 Tag and flanked by flexible GGGS linkers, was fused to the nanobodies. Biotinylation was performed as described previously(15). For the saporin-fusion, the biotinylated samples were desalted into PBS and were mixed at a concentration of 5 μM with a 2.4-fold molar excess of Streptavidin-ZAP (Advanced Targeting Systems, Carlsbad, CA, USA). The mixture was incubated for 1h on ice before diluting it in PBS to a working concentration of 500 nM.

### MicroScale Thermophoresis affinity measurements

MST affinity measurements for the binding of TCNB4, TCNB26, and TCNB34 to TC:TCblR were performed on a monolith NT.115 instrument (NanoTemper)(18). Nanobodies were labeled using the lysine reactive Monolith NT Protein Labeling kit RED-NHS (NanoTemper). As TCNB11 has a lysine residue in CDR2, it was not suitable for this labeling method and was therefore not analyzed by MST. Labeled nanobodies at concentrations ranging from 50 nM to 1.6 nM were pre-incubated with TC:TCblR at either 18 nM (for TCNB4) or 50 nM (for TCNB26 and TCNB34) for 20 min at room temperature. Subsequently, the samples were loaded into glass capillaries (NanoTemper, Munich, GER) and MST measurements were performed at various values of LED power and MST power, reaching the clearest results at 100% LED power and 80% MST power for TCNB4, at 90% LED power and 80% MST power for TCNB26 and at 90% MST power and 40% LED power for TCNB34. NanoTemper software was used to calculate the respective K_D_ values.

### Cellular nanobody uptake experiments

TCNB4 was covalently labeled with Red-NHS using the Monolith NT kit (Nanotemper technologies) and used in the cellular binding and uptake studies. HEK293 cells transfected with the *CD320* cDNA fused to a green fluorescent protein (GFP) in the protein-expression plasmid pEGFP(19). Cells were selected for stable expression of TCblR-GFP by culturing in DMEM containing G418 antibiotic and used in the present experiments. Cells were seeded at a density of 0.1 × 10^6^ cells per well in 60mm dishes, cultured for 48 h in DMEM containing 10% FBS and 200ug/ml G418.

For Fig. 2, cells were washed 3 times with PBS and incubated in DMEM with labeled TCNB4 alone or an equivalent amount preincubated for 1hr with 200ul of human serum + cobalamin as a source of holoTC. At 1, 2, or 4 h cells were washed three times with PBS and fixed in 4% paraformaldehyde for 30 min. The cell layer was washed 3 times with PBS and prepared for fluorescent microscopy. Cells incubated on ice for 1-2 h and cells incubated at 37 °C with free dye served as controls.

SI 1, cells were washed 3 times with PBS and incubated with DMEM with dsRed TC-Cbl or TCNB4+dsRed TC-Cbl. At 1, 2, or 4 h cells were washed with PBS and fixed with 4% paraformaldehyde for 30 min.

### Cell cytotoxicity Experiments

HEK293T cells in DMEM medium supplemented, with 10% v/v FBS and 200 μg/ml G418, were seeded in 96 well sterile cell culture plates at 2 × 10^3^ cells per well. After letting the cells attach for 1-2h at 37 °C. The medium was replaced with the same medium, containing 5% v/v human serum and 5 μM cyanocobalamin as a source of holoTC, as well as either 50 nM TCNB4, saporin conjugated TCNBs 4, 11, 26 or 34, unconjugated Streptavidin-ZAP as saporin control, or PBS as a negative control. After incubation for 8 days at 37 °C, the medium was exchanged to DMEM medium supplemented, with 10% v/v FBS and 200 μg/ml G418, to avoid bias by color changes in the medium and inhibition of growth respectively total cellular viability was assayed using the MTS Assay Kit (Abcam). 100% viability was defined as the readout from non-treated cells (addition of PBS) and 0% viability was defined by the readout performed on wells where no cells where seeded. Measurements were conducted in triplicates and positive and negative controls were performed in quintuplicates.

### Crystallization and data collection

After preincubating TC:TCblR at 10 mg/ml with a 1.2-fold molar excess of TCNB4 for 1h on ice, crystals of the TCNB4:TC:TCblR complex were grown by sitting drop vapor diffusion in 100 mM Bis-Tris pH 5.5, 25% w/v PEG 3350, supplemented with 10% (0.2% w/v Betaine anhydrous, 0.2% w/v L-Glutamic acid, 0.2% w/v L-Proline, 0.2% w/v Taurine, 0.2% w/v Trimethylamine N-oxide dihydrate, 0.02 M HEPES sodium pH 6.8 from the Silver Bullets Screen (Hampton Research, Viejo, CA, USA) at room temperature. The crystals were cryo-Protected by the addition of a final concentration of 25% glycerol. Diffraction data was collected from a single crystal at Swiss Light Source (SLS) at the beamline X06SA. For the data collection increments of 0.1° per 0.1 s were collected for 360° at a beam transmission of 10% and a beam size of 10 μM × 10 μM.

After preincubating TC:TCblR at 10 mg/ml with a 1.2-fold molar excess of TCNB11 for 1h on ice, crystals of the TCNB11:TC:TCblR complex were grown by sitting drop vapor diffusion in 0.2 M sodium malonate pH 6.0, 20% w/v PEG 3350. The crystals were cryo-Protected by the addition of a final concentration of 25% glycerol. Diffraction data were collected from a single crystal at Swiss Light Source (SLS) at the beamline X06SA. For the data collection increments of 0.1° per 0.1 s were collected for 360° at a beam transmission of 10% and a beam size of 10 μM × 40 μM.

After preincubating TC:CD320 at 10 mg/ml with a 1.2-fold molar excess of TCNB26 for 1h on ice, crystals of the TCNB26:TC2:CD3202 complex were grown by sitting drop vapor diffusion in 150 mM Ammonium citrate tribasic pH 7.0, 21% w/v PEG 3350. The crystals were cryo-Protected by the addition of a final concentration of 25% glycerol. Diffraction data were collected from a single crystal at Swiss Light Source (SLS) at the beamline X06SA. For the data collection increments of 0.1° per 0.1 s were collected for 360° at a beam transmission of 10% and a beam size of 20 μM × 20 μM.

After preincubating TC:TCblR at 10 mg/ml with a 1.2-fold molar excess of TCNB34 for 1h on ice, crystals of the TCNB34:TC:TCblR complex were grown by hanging drop vapor diffusion in 200 mM Ammonium Iodide, 16% w/v PEG 3350 and 1.4 × 10^5^-fold diluted seed stocks from crystals of the same protein, grown in similar conditions. The crystals were cryo-Protected by soaking in a drop containing 25% ethylene glycol. Seeds were generated using a Seed Bead™ Kit (Hampton Research, Viejo, CA, USA ). Diffraction data were collected from a single crystal at Swiss Light Source (SLS) at the beamline X06SA. For the data collection increments of 0.1° per 0.1 s were collected for 360° at a beam transmission of 10% and a beam size of 20 μM × 40 μM.

### Data processing, structure determination, model building, and refinement

Crystallographic data were indexed and reduced by XDS(20), Structures were solved by molecular replacement in Phaser MR(21) using a single monomer of TC:TCblR (PDB ID 4ZRP)(15) as input model. Nanobodies were fitted based on an existing model of a nanobody (PDB ID 3P0G)(22). Model building was done in Coot(23) and Refinement was performed in Phenix(24). To avoid bias from the comparably low-resolution map of TCNB26:TC_2_-TCblR, for this structure only rigid body and B-factor refinement was performed.

### Figure preparation and data analysis

Protein sequence alignments were performed in CLC Genomics Workbench. Cell viability assay data were analyzed and plotted in GraphPad Prism. Protein structure images were generated in PyMol(25).

## Results

### Generation of nanobodies against TC:TCblR complex

We immunized an alpaca with the purified, B_12_-bound complex of TC and TCblR (in the following named TC:TCblR)(15) and selected 20 unique binders (nanobodies) by phage display(17). Fourteen binders formed stable complexes with TC:TCblR, as confirmed by size exclusion chromatography (SEC). Four of these nanobodies (TCNB4, TCNB11, TCNB26, TCNB34) formed well-diffracting crystals in complex with TC:TCblR (Fig, 1a). The affinity of nanobodies TCNB4, TCNB26 and TCNB34 to the TC: TCblR complex was determined by MicroScale thermophoresis (MST)(18), as they were compatible with the required amine-reactive labelling(26). This revealed dissociation constants (KD) of 5.2 nM for TCNB4, 87.4 nM for TCNB26, and 12.8 nM for TCNB34 (Fig. 1b).

**Figure 1.**
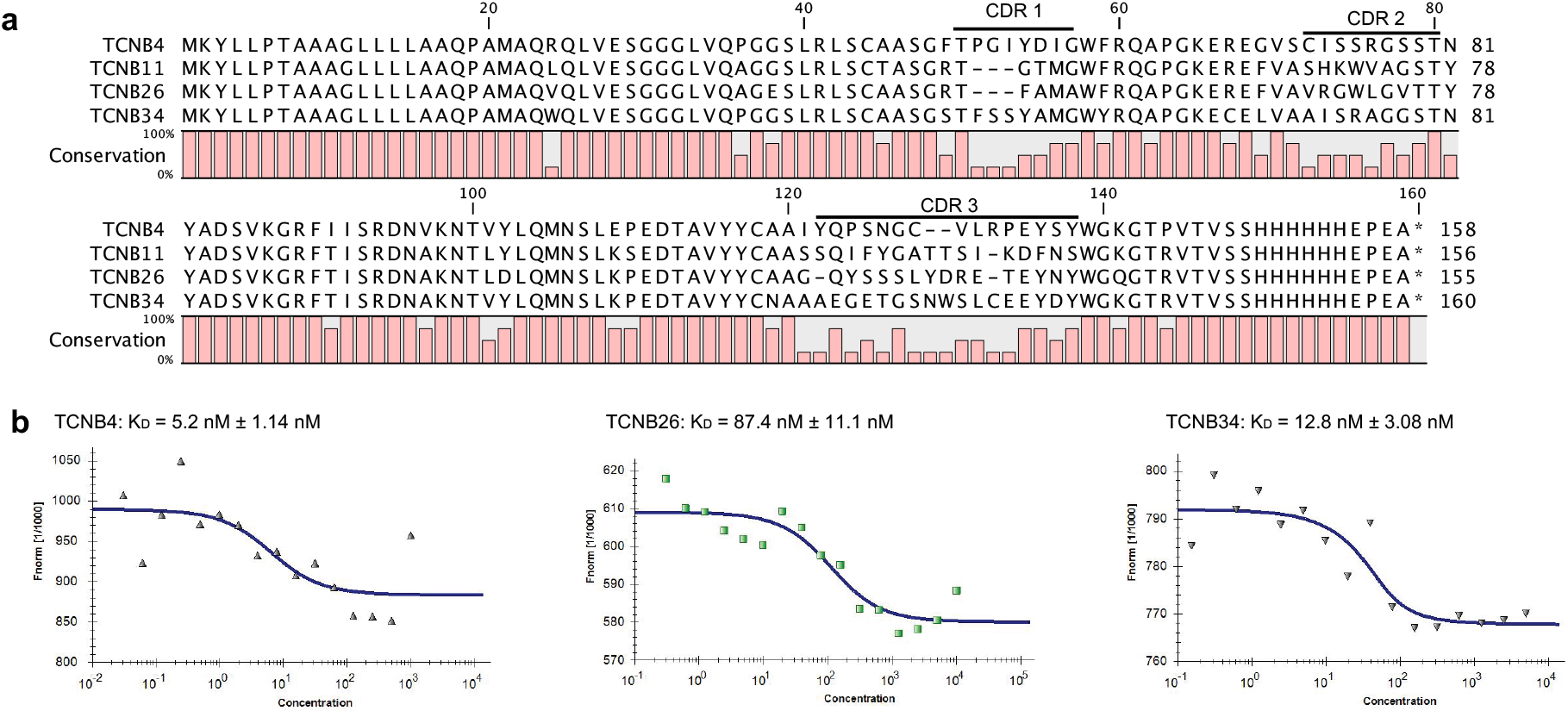
Biophysical characterization of nanobodies raised against TC:TCblR. **a**, Sequence alignment of selected conformationally binding nanobodies. Red bars below the sequence indicate the degree of sequence conservation. **b**, MicroScale thermophoresis affinity measurements of nanobodies TCNB4, TCNB26, and TCNB34 binding to TC:TCblR.

### Cellular uptake of TCNB4

We selected the highest affinity binder, TCNB4, to test its potential for TC:TCblR-mediated cellular uptake. We incubated fluorescently labeled TCNB4 with HEK293 cells that constitutively overexpressed GFP-tagged TCblR (GFP-TCblR)(19). Using light microscopy, we observed co-localization of TCNB4 and TCblR (Fig 2). We further observed that the nanobody and the receptor protein were endocytosed. To test whether endocytosed nanobody was degraded, we repeated the experiment with cells overexpressing a fusion construct of TC and red fluorescent protein (DsRed)(27). Over the course of four hours, we observed the loss of DsRed fluorescence, indicating lysosomal degradation of DsRed-TC. For the labeled TCNB4 however, we did not observe any loss of the fluorescence signal, indicating that the nanobody remained intact upon endocytic internalization (SI Figure 1). Given that the fluorophore has an extended life span, some degradation of the nanobody cannot be excluded.

**Figure 2.**
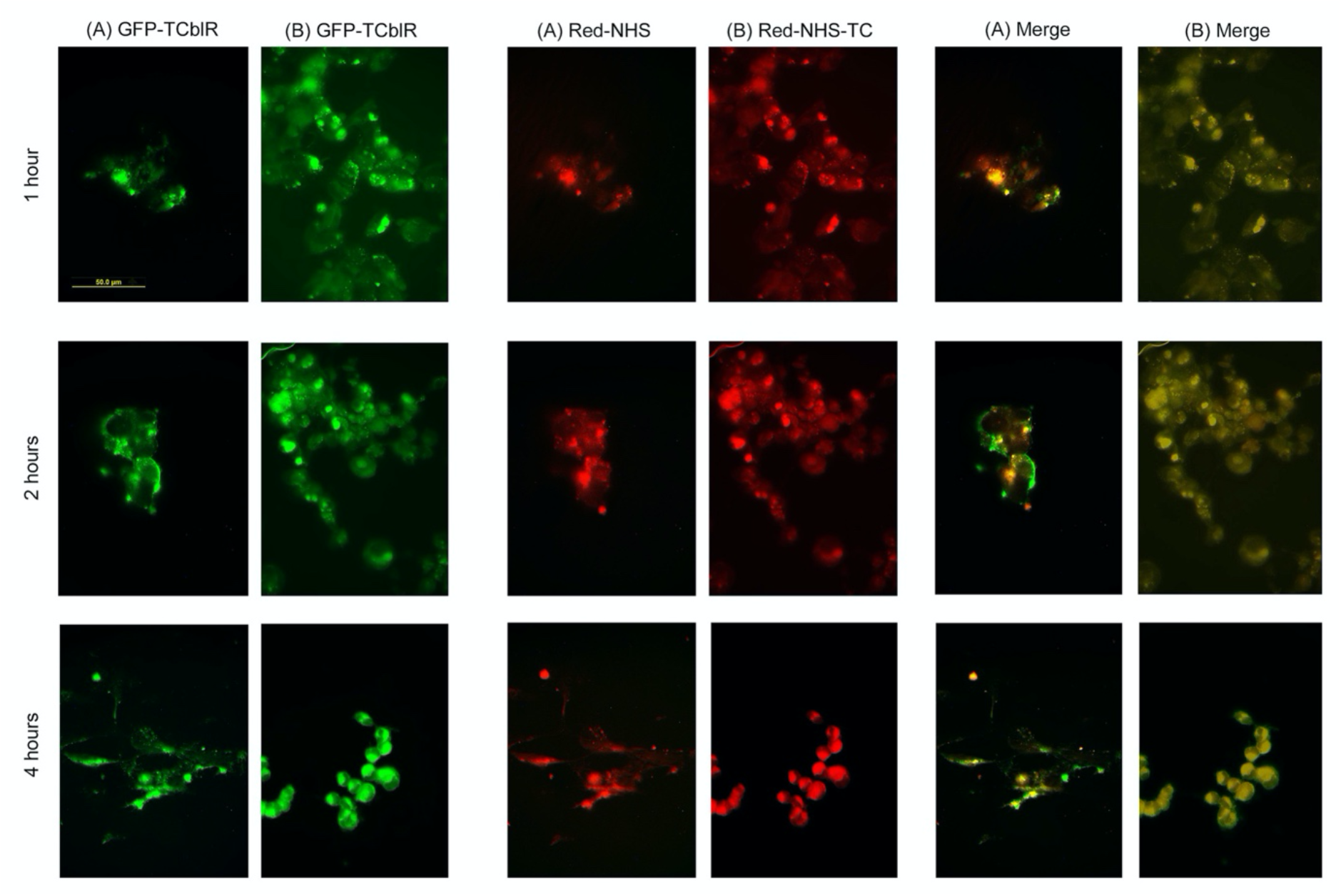
Nanobody uptake by HEK 293 cells expressing TCblR-GFP. Cells were incubated with red fluorescence labeled TCNB4 (A) or with labeled TCNB4 + TC-Cbl (B) for 1 h, 2 h, or 4 h at 37 °C. Over the time course tested, TCNB4 in the absence of TC appears to bind to some extent to the cell surface, which may be due to direct (and low affinity) interaction with the extracellular domain of TCblR (panels A), but without internalization (the merge of panels A). Nanobody pre incubated with TC-Cbl appears to bind specifically to CD320-GFP (panels B) and is associated with CD320-GFP internalization (merge panels B). However, the CD320-GFP and the red nanobody do not appear to degrade at 4h as seen by intense fluorescence of TCblR-GFP and the nanobody.

### Inhibition of cell growth by saporin-conjugated nanobodies

To investigate whether nanobodies carrying toxic cargo could kill HEK293T cells we fused them to the cytotoxic, plant-derived enzyme saporin, which acts by irreversibly inactivating eukaryotic ribosomes(28). The fusion was accomplished by inserting a biotinylation site (AVI-tag) into the nanobodies and biotinylating them using the enzyme BirA(29). The biotinylated nanobodies were then coupled with biotin-reactive saporin (Streptavidin-ZAP, Advanced Targeting systems). The resulting nanobody-saporin conjugates were used for cytotoxicity assays.

We seeded highly proliferating HEK293T cells at low density to stimulate the expression of *CD320*(8). When adding the saporin-conjugated nanobodies at a concentration of 50 nM, complete inhibition of cell growth was observed for TCNB4 (Fig. 3a). This effect was not observed for unconjugated TCNB4 or unconjugated saporin-ZAP (Fig. 3b). saporin-conjugated TCNB11 and TCNB34 also inhibited cell growth, but to a lower extent than TCNB4. Interestingly, saporin-conjugated TCNB26 did not show significant cytotoxicity. This might be due to the lower affinity of TCNB26 compared to the other nanobodies.

**Figure 3.**
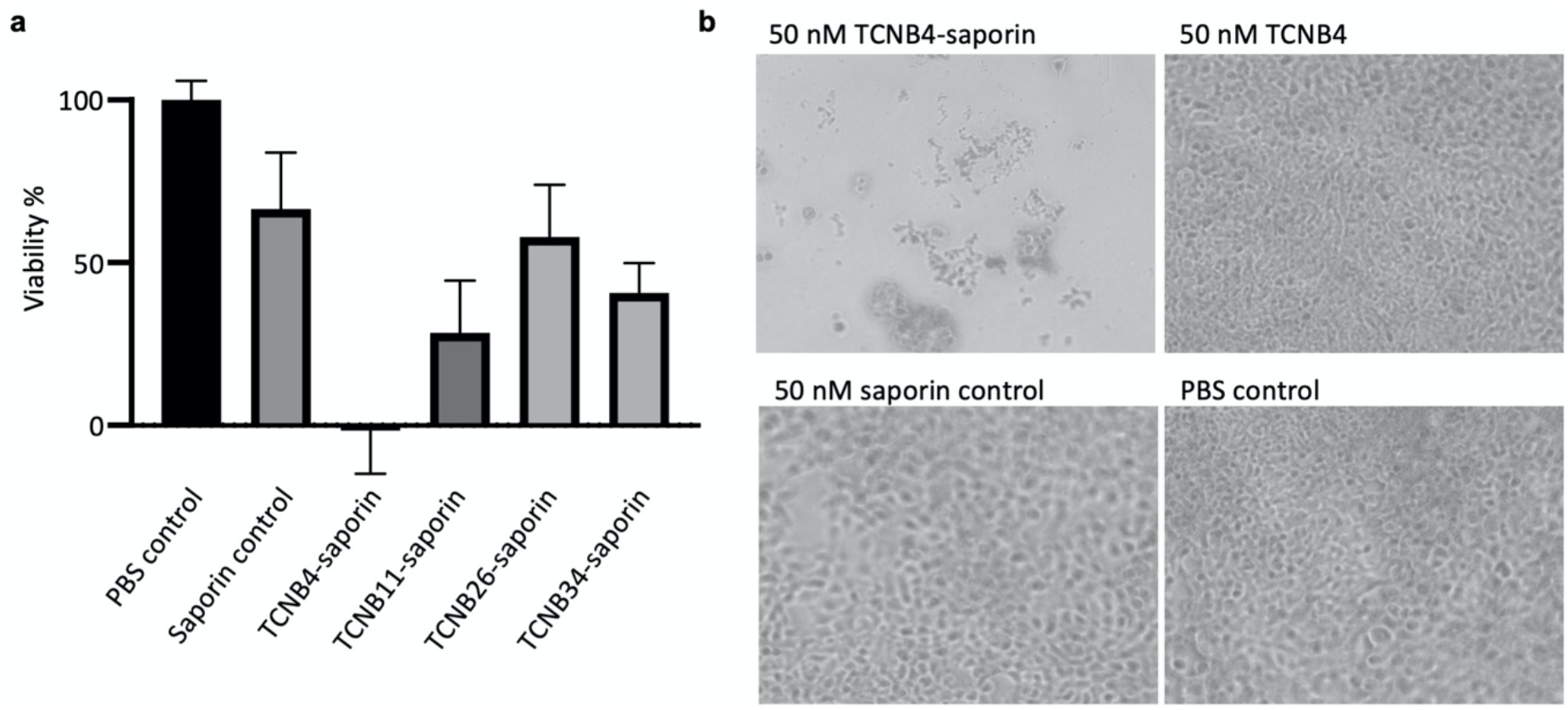
Cytotoxicity assay for saporin-conjugated nanobodies. **a**, The viability of HEK293T cells after incubation with saporin-conjugated nanobodies for 8 days was determined by measuring the conversion of MTS tetrazolium reagent by viable mammalian cells, which generates a colored formazan dye. The viability of untreated cells (PBS buffer added) was set to 100%, whereas dye conversion in absence of any cells was set to 0%. **b**, Light microscopy images of HEK293T cells after incubation with saporin-conjugated TCNB4, TCNB4 alone, saporin alone, or PBS buffer (control) for 8 days.

### Structural characterization of nanobody binding

To delineate the interaction interfaces, we determined individual crystal structures of all four nanobodies in complex with TC:TCblR (Fig. 4, SI Table 1). The structures revealed three distinct binding epitopes. TCNB11, TCNB26, and TCNB34 bind to TC only, whereby TCNB11 and TCNB34 share overlapping binding epitopes. In contrast, TCNB4 binds at the interface of TC and TCblR and therefore specifically recognizes the TC:TCblR complex.

**Figure 4.**
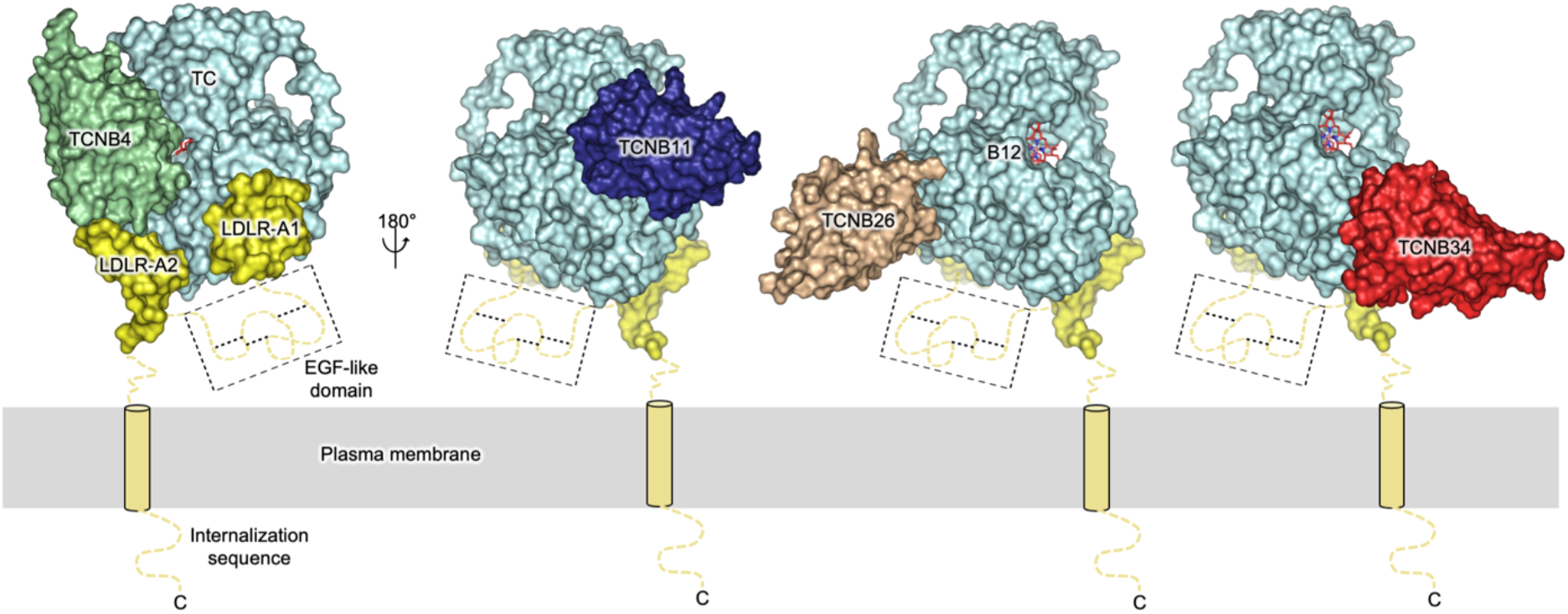
Structural epitopes on TC:TCblR. Surface representation of the crystal structures of TC:TCblR:nanobody complexes with TC colored cyan, the LDLR-A1 and LDLR-A2 domains of CD320 colored yellow, and vitamin B_12_ in red stick representation. Flexible regions of TCblR that were not resolved in the crystal structures (EGF-like domain) or were truncated in the crystallization construct (N-terminus including transmembrane helix) are depicted schematically. Nanobodies are shown as surfaces colored green (TCNB4), blue (TCNB11), brown (TCNB26), and red (TCNB34).

TCNB4 interacts with TCblR via complementarity-determining region 1 (CDR1) and a non-variable loop (Fig. 4, SI Figure 2). CDR3 mediates most interactions with TC. A disulfide bond between CDR2 and CDR3 restricts the conformational freedom of CDR3 and presumably helps with its precise positioning. We observed density likely reflecting a Ca^2+^ ion coordinated by backbone carbonyl groups of CDR3 and a glutamate side chain of CDR1. The coordinated ion appears to stabilize the conformation of the CDR3 loop further. TCNB11 binds its TC by means of contacts involving CDR2 and CDR3 (Fig. 4, SI Figure 3). CDR3 twists a flexible loop of TC near His173 by 180°. The flexibility of this loop has been observed previously: While His173 replaces the upper coaxial ligand of hydroxocobalamin(30), the cyano-group cyanocobalamin stays bound and displaces His173 from the B_12_ binding site(15). TCNB26 binds to TC with all three CDRs (Fig. 4, SI Figure 4). The relatively loosely packed binding interface might explain the comparably lower binding affinity of this nanobody. TCNB34 binds TC predominantly via CDR1 and CDR3 (Fig. 4d, SI Figure 5). CDR2 forms a disulfide bond to CDR3, similar to TCNB4.

### Novel structural insight into the TC:TCblR complex

The crystal structure of TC:TCblR bound to TCNB34 improved the previously reported resolution of 2.1 Å(15) to 1.85 Å (SI Table 1). While the resolution was higher overall, two regions in particular were better resolved (Fig. 5). First, we were able to build a complete model of the interface of TC and the LDLR-A1 domain of TCblR. In the previously published structures of TC, either the loop connecting Cys65 and Cys78 was disordered(15) or the disulfide bond between the two cysteines was not formed(30). In our TC:TCblR:TCNB34 structure, the disulfide bond is formed and the loop is well resolved, revealing a different conformation than previously observed. Moreover, the structure reveals new binding interactions between TC and TCblR. Glu73 and Asp74 of TC form salt bridges with Lys58 of TCblR. As salt bridges are sensitive to changes in pH(31), this observation can help rationalize the pH-dependent dissociation of TC from TCblR in the lysosome after endocytosis(15). We also observed a network of ordered water molecules that stabilize the complex of TC and TCblR by H-bonding. Second, our high-resolution map provided a more detailed view of the region around the disease-related residue Glu88 in CD320(32). Deletion of this residue has been associated with vitamin B_12_ deficiency(32). Our structure shows that the adjacent residue Glu87 forms two salt bridges: One with Arg68 via its carboxyl group and one with Arg73 via its backbone carbonyl. The newly revealed salt bridges indicate a possible role of Glu88 in pH-dependent cargo release of the receptor upon endocytosis. This region is also involved in Ca^2+^ binding. Thus, genetic deletion of Glu86, Glu87, or Glu88 would lead to the loss of the observed salt bridges, thereby destabilizing the receptor structure. This finding is in line with the lower affinity of the ΔGlu88 mutant receptor to TC(15).

**Figure 5.**
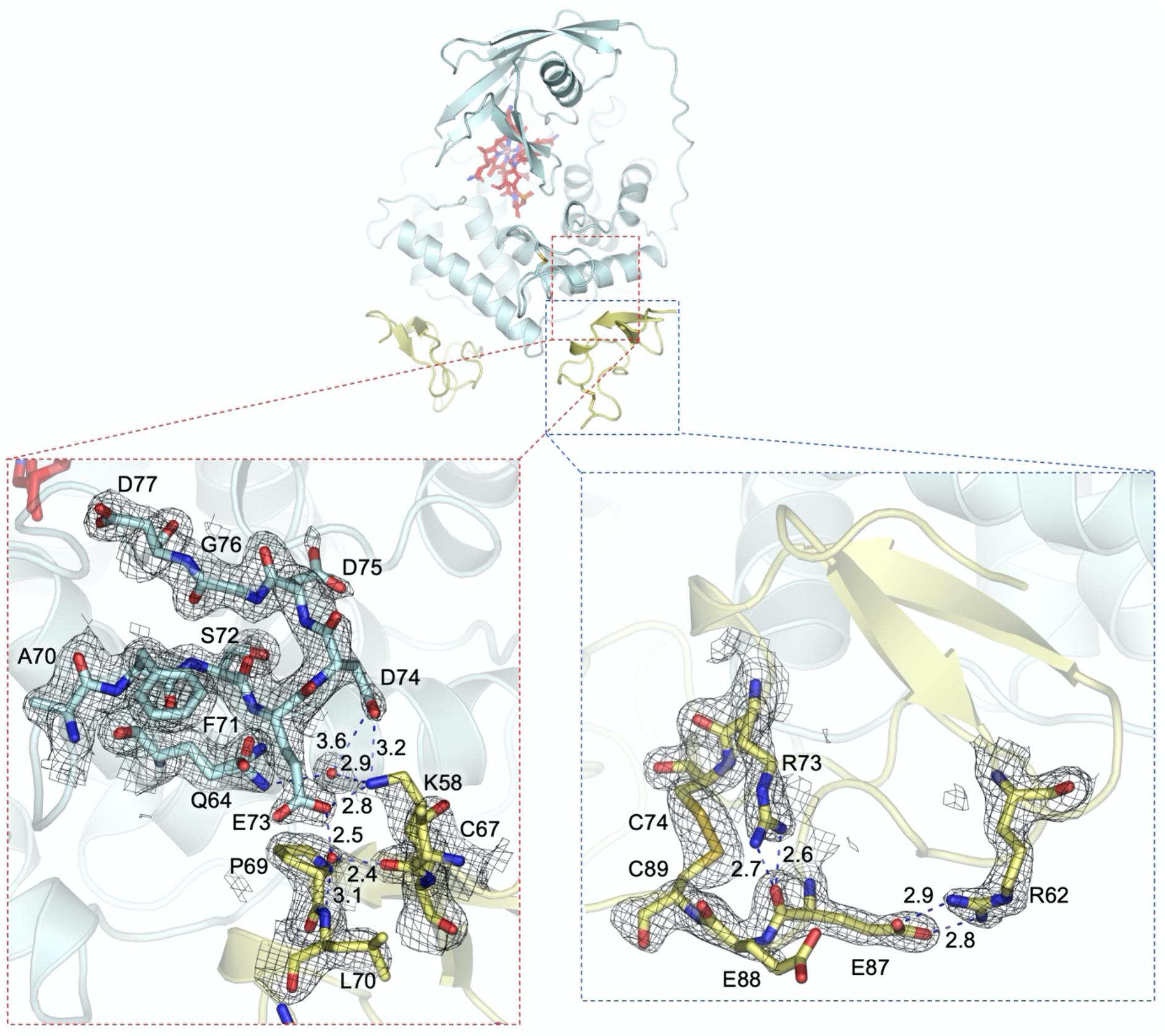
Structural insights into transcobalamin recognition by its receptor TCblR. Two close up views of the interaction interface between TC and TCblR as revealed in the co-crystal structure of TC:TCblR:TCNB34 are shown. (dashed lines). TC (colored cyan) and TCblR (colored yellow) are shown in cartoon representation, TCNB34 is omitted for clarity. Bound B12 (colored red) and disulfide bonds forming the loops of interest are shown in stick representation. The left inset shows a loop in TC that was disordered in previous crystal structures(15, 30). Polar contacts between TC and TCblR are indicated with dashed blue lines, with distances indicated in Å. The right inset shows the TCblR residue Glu88 and its surrounding. Salt bridges involving Glu87 are indicated with blue dashed lines. A genomic deletion of Glu88, which would be equal to deletion of Glu87, was reported to lead to B12 deficiency(32). The black mesh represents a 2Fo-Fc map contoured at 1.0 σ and carved to 1.6 Å.

The TCNB11:TC:TCblR co-crystal structure reveals the structure and position of eight residues connecting LDLR-A1 and the EGF-like domain, including Cys94. As Cys94 is already part of the EGF-like domain, this finding may be indicative of the location of the otherwise disordered EGF-like domain (SI Figure 3c).

## Discussion

We have generated four high-affinity nanobodies targeting TC or the TC:TCblR complex. Given that these nanobodies can enter human cells and inhibit cell growth when fused to a toxin, they may have a value in diagnostic or therapeutic approaches. As the nanobodies bind distinct epitopes, fusing two of them (for example, TCNB4 and TBNB11) might increase the affinity or avidity of TC:TCblR:nanobody complex formation *in vivo*. One might expect that such a bipartite nanobody could bind TC in the blood before binding to TCblR. To determine the applicability of our nanobody-drug conjugates, further studies *in vivo* in native environments will be required. Future investigations may also test the fusion of our nanobodies to other toxins or drugs for cancer therapy or radionuclides for tumor localization using imaging studies.

Our crystal structures revealed new features of the TC:CD320 complex, which helped further rationalize the lysosomal dissociation of TC-Cbl from TCblR at low pH(15) as well as the structural basis of vitamin B_12_ deficiency related to the ΔE88 mutation(32). The structural role of the EGF-like domain in TCblR remains elusive. Future endeavors employing methods such as 3D variability analysis of cryo-electron microscopy data(33) may be key to revealing the role of the inherently mobile domain.

## Nonstandard abbreviations

*CD320*: Gene of transcobalamin receptor protein
CD320/TCblR: Trascobalamin receptor protein
DMEM: Dulbecco’s Modified Eagle Medium
GFP: Green fluorescent protein
HEPES: 4-(2-hydroxyethyl)-1-piperazineethanesulfonic acid
MST: MicroScale Thermophoresis
PBS: Phosphate buffered saline
TC: Transcobalamin II

## Acknowledgments

We thank the beamline staff at the Swiss Light Source at the Paul Scherrer Institute in Villigen for assistance with data collection. We thank Dr. S. Stefanic (University of Zurich) for help with alpaca immunizations, blood collection, and testing of the specific immune response by ELISA. We thank Prof. R. Dutzler (University of Zurich) for technical support and the primers and vectors used for the generation of the VHH DNA library. We thank M. Mikolin for help with cell culture maintenance. This work was supported by the Swiss National Science Foundation (SNF 310030B_166672 to K.P.L.).

## Author contributions

J.S.B. and K.P.L conceived the project. J.S.B. designed and performed all experiments, except for the cellular uptake experiments with fluorescently labeled TCNB4 which were performed by E.Q and J.M.S., and generation of the VHH library which was performed by A.S.R., J.S.B. and K.P.L. analyzed the data and wrote the paper. All authors contributed to the final version of the manuscript.

## Supplementary information

**SI Figure 1.**
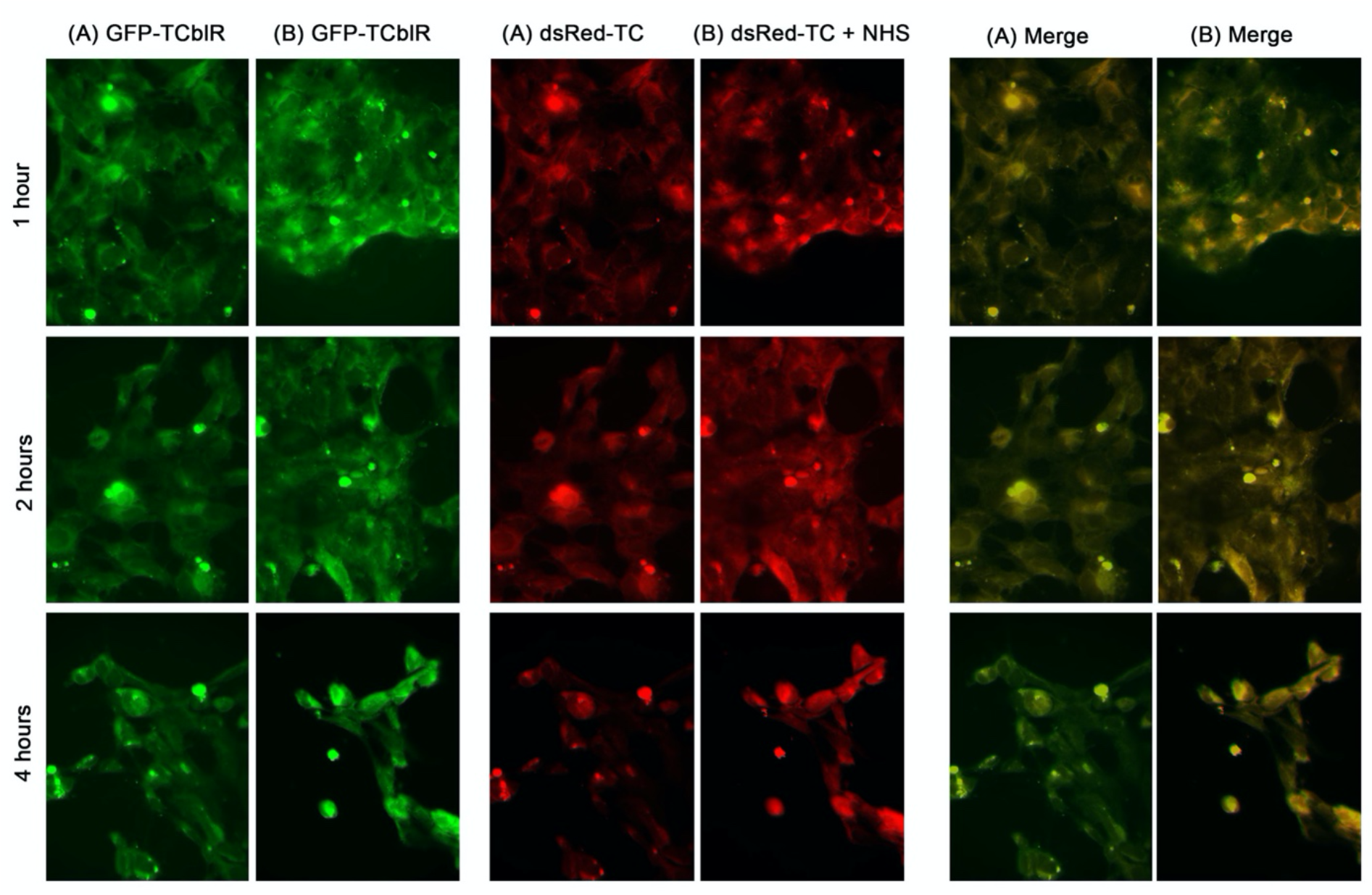
Internalization of dsRed TC-Cbl and dsRed TC-Cbl–TCNB4 by HEK293 cells expressing TCblR-GFP. Cells were incubated with dsRedTC-Cbl (A) or with nanobody TCNB4 +dsRed TC-Cbl for 1, 2, or 4 h at 37 °C. Over time, the dsRed-TC-Cbl is internalized and is degraded (panels A). However, the TCNB4 + dsRed TC-Cbl is internalized but is not degraded (panels B).

**SI Figure 2.**
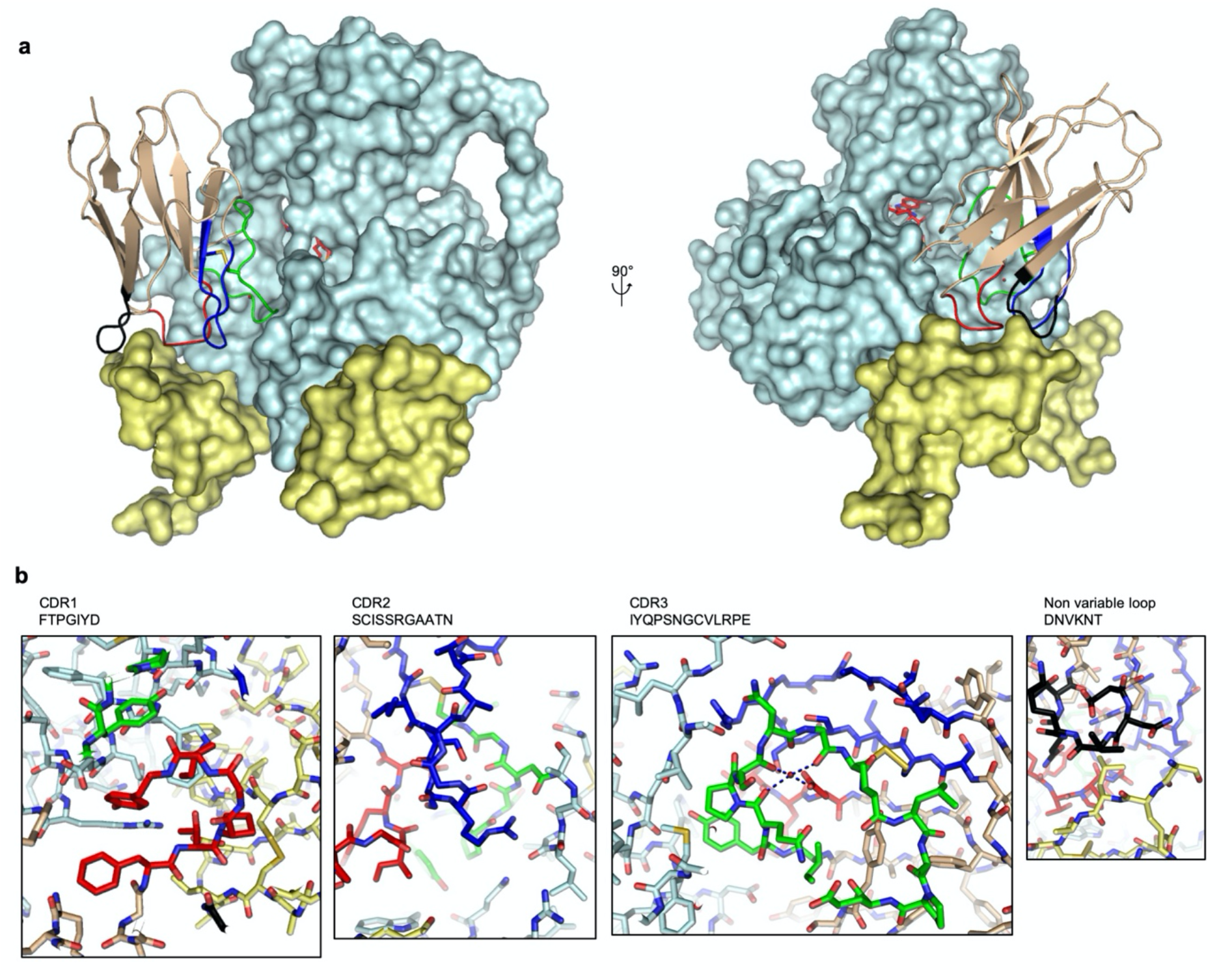
Crystal structure of TC:TCblR bound to TCNB4. **a**, TC (cyan) and TCblR (yellow) are shown in surface representation. Cyanocobalamin (red) is shown in stick representation. TCNB4 (brown) is shown in cartoon representation with CDR1 (red), CDR2 (blue), CDR3 (green), as well as a non-variable loop (black) that are interacting with TCblR are highlighted with colors. A disulfide bond between CDR2 and CRD3 is shown in stick representation. Note that the nanobody is interacting with both TC as well as the receptor TCblR. **b**, Close upview on molecular interaction between different CDR loops of the nanobody and TC:TCblR shown in stick representation, using the same colors as in a. Note that in CDR3, there seems to be a Ca^2+^ ion coordinated by three backbone carboxyl groups of CDR3 and by one aspartate of CDR1.

**SI Figure 3.**
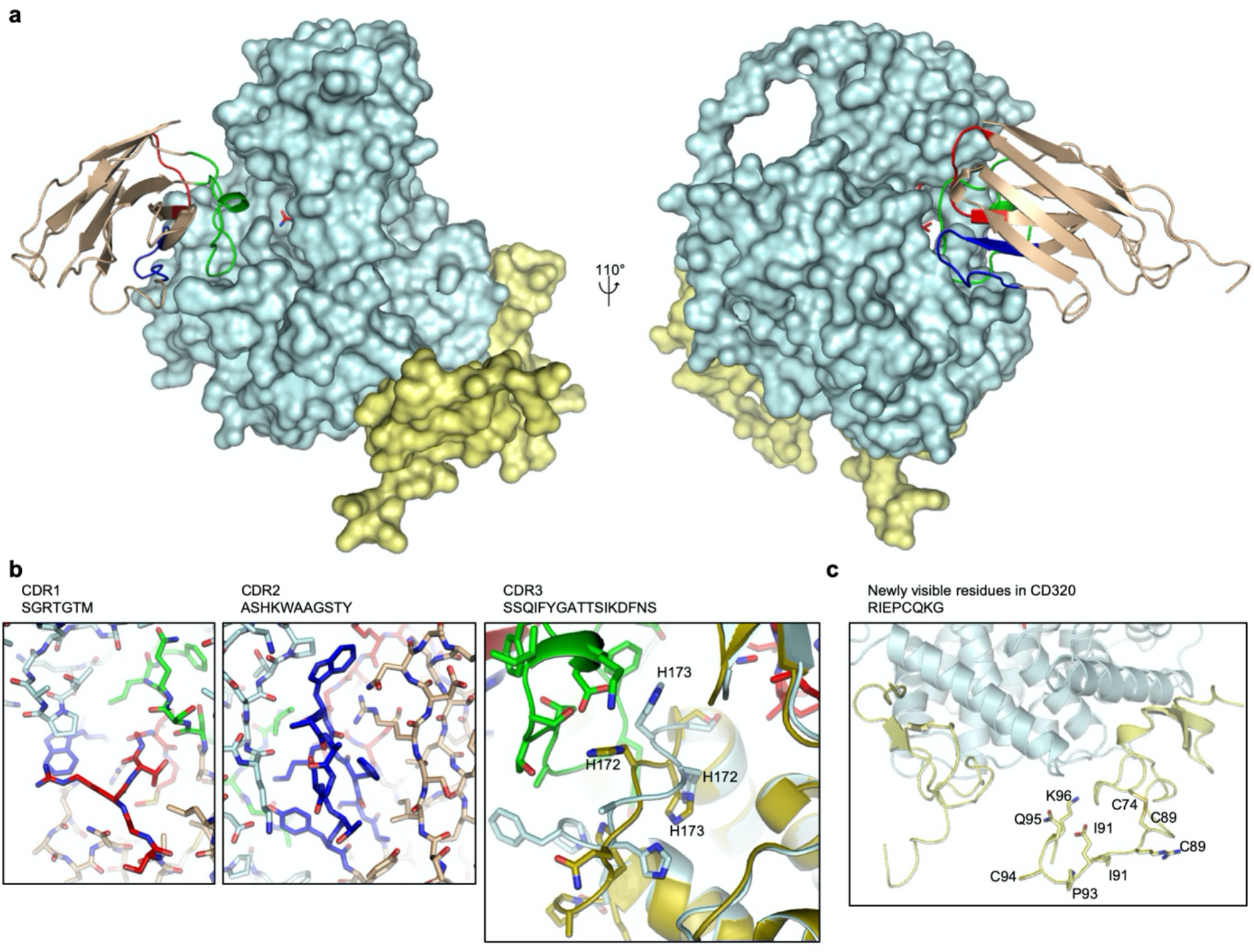
Crystal structure of TC:TCblR bound to TCNB11. **a,** TC (cyan) and TCblR (yellow) are shown in surface representation. Cyanocobalamin (red) is shown in stick representation. TCNB11 (brown) is shown in cartoon representation with CDR1 (red), CDR2 (blue), CDR3 (green) are highlighted with colors. **b**, Close-up view on molecular interaction between different CDR loops of the nanobody and TC shown in stick representation, using the same colors as in a. In the panel, for CDR3, the structure of TC has been superposed with a nanobody free structure of TC (gold) (4ZRP(15)) and for clarity, the structures are shown in cartoon representation, and only selected residues and B12 are shown in stick representation. Note that the binding of TCNB11 induces a twist in TC in a loop at the binding interface. **c**, Residues of TCblR extending LDLR-A1 that were not resolved in previous structures, as well as the disulfide-bond forming residues Cys74 and Cys89 are shown in stick representation.

**SI Figure 4.**
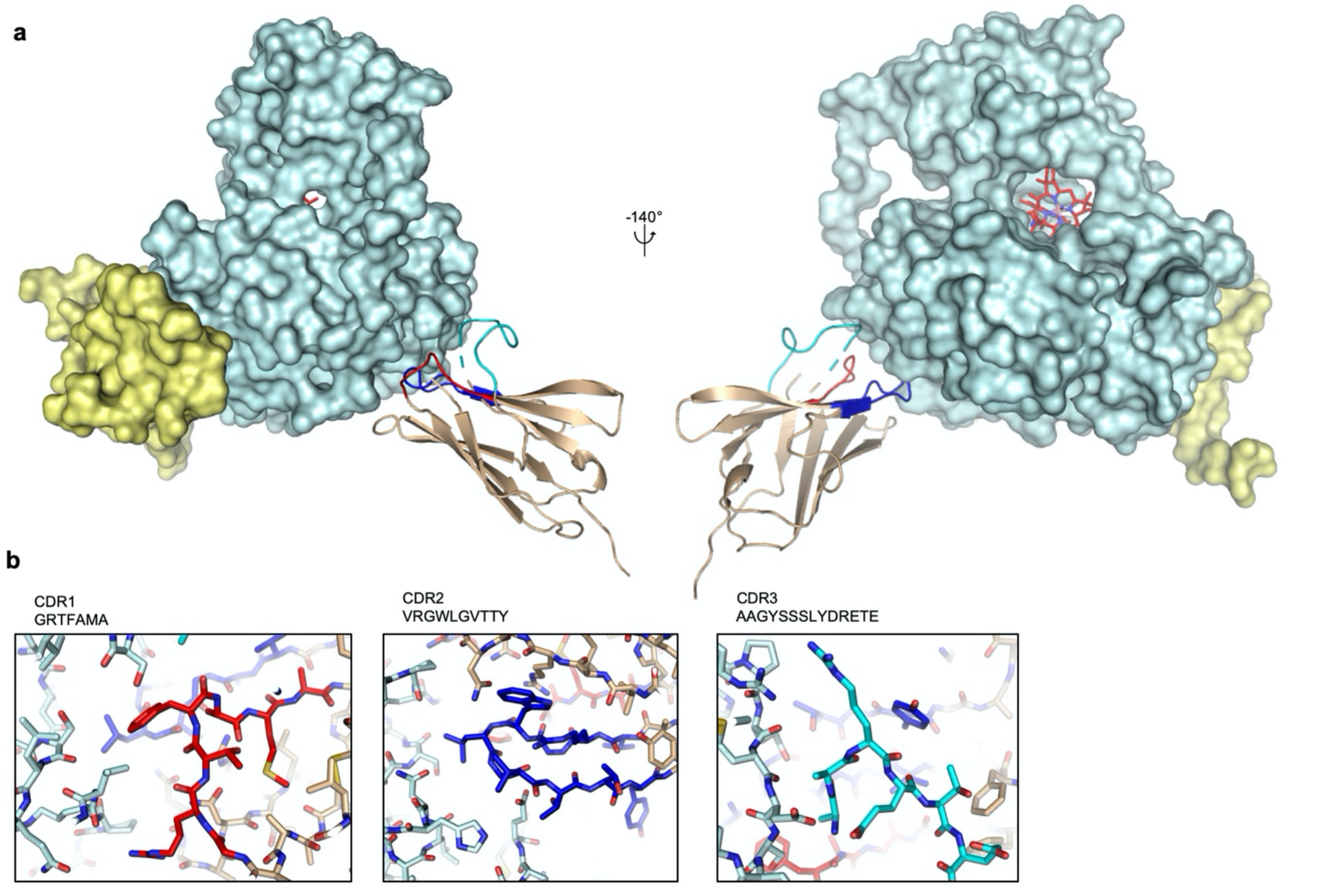
Crystal structure of TC:TCblR bound to TCNB26. **a**, TC (cyan) and TCblR (yellow) are shown in surface representation. Cyanocobalamin (red) is shown in stick representation. TCNB26 (brown) is shown in cartoon representation with CDR1 (red), CDR2 (blue), CDR3 (green) highlighted with colors. **b**, Close up view on molecular interaction between different CDR loops of the nanobody and TC, shown in stick representation, using the same colors as in **a**.

**SI Figure 5.**
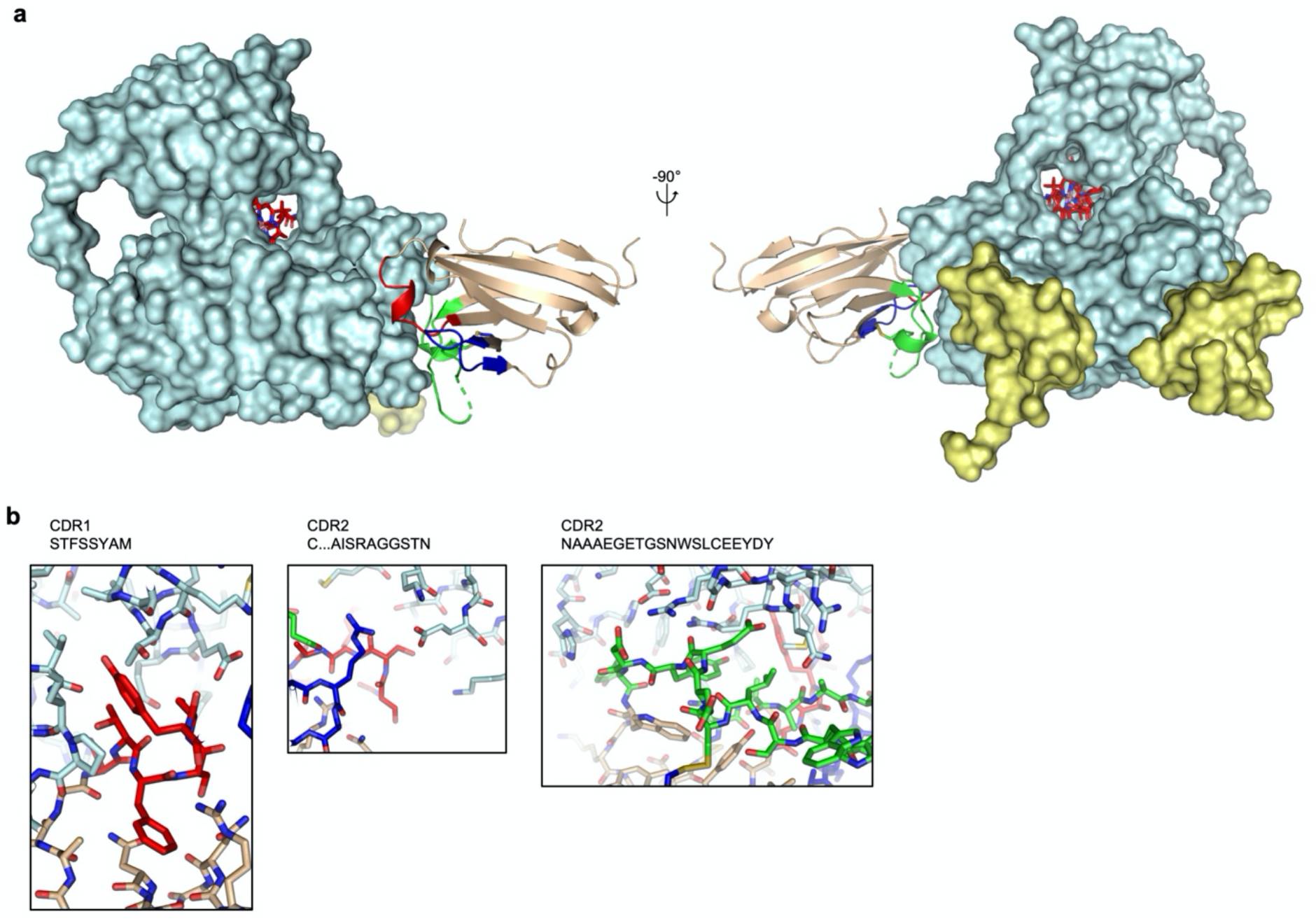
Crystal structure of TC:TCblR bound to TCNB34. **a**, TC (cyan) and TCBlR (yellow) are shown in surface representation. Cyanocobalamin (red) is shown in stick representation. TCNB34 (brown) is shown in cartoon representation with CDR1 (red), CDR2 (blue), CDR3 (green) highlighted with colors. A disulfide bond between CDR2 and CRD3 is shown in stick representation. **b**, Close-up view on molecular interaction between different CDR loops of the nanobody and TC shown in stick representation, using the same colors as in **a.**

**SI Table 1.**

|  | TCNB4:TC:CD320 | TCNB11:TC:CD320 | TCNB26:TC<sub>2</sub>:CD320<sub>2</sub> | TCNB34:TC:CD320 |
| --- | --- | --- | --- | --- |
| Wavelength | 0.979502 | 1 | 0.979502 | 0.979502 |
| Resolution range | 48.92 - 2.692 (2.789 - 2.692) | 48.96 - 3.0 (3.107 - 3.0) | 49.14 - 4.184 (4.334 - 4.184) | 49.27 - 1.854 (1.92 - 1.854) |
| Space group | P 21 21 21 | C 2 2 21 | P 43 21 2 | P 43 21 2 |
| Unit cell | 105.339 116.418 125.498 90 90 90 | 103.717 195.831 198.385 90 90 90 | 102.129 102.129 360.998 90 90 90 | 104.822 104.822 167.46 90 90 90 |
| Total reflections | 84243 (7177) | 81479 (8030) | 28382 (1406) | 158598 (15319) |
| Unique reflections | 42564 (3745) | 40763 (4011) | 14241 (752) | 79459 (7693) |
| Multiplicity | 2.0 (1.9) | 2.0 (2.0) | 2.0 (1.9) | 2.0 (2.0) |
| Completeness (%) | 97.98 (87.42) | 99.84 (99.80) | 94.99 (51.80) | 99.79 (98.15) |
| Mean I/sigma(I) | 7.97 (1.17) | 11.45 (2.30) | 10.17 (1.59) | 19.37 (1.89) |
| Wilson B-factor | 56.39 | 68.96 | 164.21 | 31.94 |
| R-merge | 0.08061 (0.6841) | 0.05523 (0.371) | 0.04472 (0.4819) | 0.0272 (0.4422) |
| R-meas | 0.114 (0.9675) | 0.07811 (0.5247) | 0.06324 (0.6815) | 0.03847 (0.6254) |
| R-pim | 0.08061 (0.6841) | 0.05523 (0.371) | 0.04472 (0.4819) | 0.0272 (0.4422) |
| CC1/2 | 0.993 (0.459) | 0.997 (0.791) | 0.999 (0.851) | 0.999 (0.66) |
| CC* | 0.998 (0.793) | 0.999 (0.94) | 1 (0.959) | 1 (0.892) |
| Reflections used in refinement | 42558 (3745) | 40739 (4008) | 14207 (748) | 79457 (7693) |
| Reflections used for R-free | 2128 (187) | 2037 (201) | 712 (37) | 3973 (384) |
| R-work | 0.2287 (0.3664) | 0.2057 (0.3629) | 0.2666 (0.3837) | 0.1731 (0.2875) |
| R-free | 0.3015 (0.3973) | 0.2731 (0.4116) | 0.2866 (0.3419) | 0.2044 (0.3046) |
| CC(work) | 0.933 (0.671) | 0.948 (0.792) | 0.843 (0.681) | 0.957 (0.824) |
| CC(free) | 0.896 (0.494) | 0.895 (0.695) | 0.694 (0.644) | 0.938 (0.784) |
| Number of non-hydrogen atoms | 9598 | 9548 | 8372 | 5545 |
| macromolecules | 9352 | 9354 | 8182 | 4826 |
| ligands | 193 | 191 | 190 | 312 |
| solvent | 53 | 3 | 1070 | 407 |
| Protein residues | 1217 | 1217 | 0.008 | 612 |
| RMS(bonds) | 0.016 | 0.018 | 1.24 | 0.012 |
| RMS(angles) | 1.73 | 1.75 | 93.89 | 2.58 |
| Ramachandran favored (%) | 86.11 | 84.04 | 5.15 | 98.17 |
| Ramachandran allowed (%) | 10.29 | 10.36 | 0.95 | 1.83 |
| Ramachandran outliers (%) | 3.6 | 5.6 | 1.36 | 0 |
| Rotamer outliers (%) | 0.1 | 0 | 8.9 | 0.76 |
| Clashscore | 29.25 | 20.96 | 213.36 | 9.14 |
| Average B-factor | 74.07 | 74.18 | 214.3 | 40.38 |
| macromolecules | 74.34 | 74.33 | 172.98 | 40.28 |
| ligands | 66.01 | 66.99 |  | 34.87 |
| solvent | 54.3 | 57.6 |  | 45.8 |

